# Cannabinoid inhibition of mechanosensitive K^+^ channels

**DOI:** 10.1101/2024.12.09.627564

**Authors:** Trevor Docter, Ben Sorum, Rahul Deshmane, Cody Doubravsky, Stephen G. Brohawn

**Author notes:** Department of Biomedical Sciences, Cooper Medical School of Rowan University, Camden, NJ, 08103, USA.

## Abstract

Cannabidiol (CBD) is a prominent non-psychoactive small molecule produced by cannabis plants used clinically as an antiepileptic. Here, we show CBD and other cannabinoids are potent inhibitors of mechanosensitive two-pore domain K^+^ (K2P) channels, including TRAAK and TREK-1 that contribute to spike propagation in myelinated axons. Five TRAAK mutations that cause epilepsy or the neurodevelopmental syndrome FHEIG (facial dysmorphism, hypertrichosis, epilepsy, intellectual/developmental delay, and gingival overgrowth) retain sensitivity to cannabinoid inhibition. A cryo-EM structure reveals CBD binds in the intracellular cavity of TREK-1 to sterically block ion conduction. These results show that cannabinoids and endogenous lipids compete for a common binding site to inhibit channel activity, identify mechanosensitive K2Ps as potential physiological targets of CBD, and suggest cannabinoids could counter gain-of-function in TRAAK channelopathies.

**Summary:** We discover cannabinoids inhibit mechanosensitive K+ channels including mutants that cause disease and reveal the mechanism for channel block.

## Introduction

Phytocannabinoids are hydrophobic phenolic compounds produced by cannabis plants. Over one hundred phytocannabinoids have been identified, with delta-9-tetrahydrocannabinol (THC) and cannabidiol (CBD) among the most prevalent ^1,2^. THC is the primary psychoactive cannabinoid that underlies the euphoria-inducing effects of cannabis ^1^. The effects of THC are mainly mediated by partial agonism of CB1 and CB2 G-protein coupled receptors (EC_50_ ~ 50nM) ^1^. CBD is the predominant non-psychoactive cannabinoid and is an effective antiepileptic approved by regulatory agencies in multiple jurisdictions for the treatment of seizures in Lennox-Gastaut syndrome, Dravet’s syndrome, and tuberous sclerosis complex ^3–8^. In contrast to THC, the molecular basis for the antiepileptic effects of CBD is unknown ^1,2^. CBD is not a CB1 or CB2 receptor agonist; rather, it negatively modulates CB1 and GPR55 receptors (IC_50_ ~500 nM)^9–11^. Like other classic anticonvulsants, CBD inhibits voltage-gated sodium channels (IC_50_ ~2-10 µM) ^12,13^. However, the relevance of these targets is debated^2^, as CBD’s antiepileptic effects are observed in slice recordings at concentrations ≤100 nM and in treated patients at ~50-100 nM (as measured in plasma or estimated in brain tissue). CBD also modulates other ion channels, activating TRPV1-4 (EC_50_ ~1-4 µM), TRPA1 (EC_50_ 110 nM) ^14,15^, and Kv7.2/7.3 (EC_50_ 200 nM) ^16,17^, and inhibiting BK (IC_50_ 280 nM) ^18^, but the importance of these channels to CBD pharmacology is similarly unverified.

TRAAK (TWIK-related arachidonic acid-activated K+ channel) is a mechanosensitive potassium-selective ion channel belonging to the two-pore domain (K2P) K^+^ ion channel family. TRAAK is localized to axon initial segments ^19,20^ and nodes of Ranvier ^21,22^, where it controls the resting membrane potential and facilitates action potential repolarization, thereby enabling high frequency spiking. Gain-of-function TRAAK mutations cause epileptic and neurodevelopmental disorders in humans, with four reported variants in FHEIG (facial dimorphism, hypertrichosis, epilepsy, intellectual disability, and gingival outgrowth) syndrome and one in Rolandic epilepsy ^23–28^.

TRAAK displays low leak activity under resting conditions and is highly activated by increasing membrane tension^29,30^. Conformational equilibrium between up and down states of transmembrane helix 4 (TM4) from each subunit of the dimeric channel underlies mechanical activation ^30–32^. At low tension, the TM4 down state predominates, creating membrane-facing fenestrations above TM4 that connect the channel cavity to the surrounding membrane inner leaflet. In this conformation, lipid acyl chains access the channel cavity through the open fenestrations to block conduction. Leak activity is due to rare spontaneous delipidation of the TM4 down state, resulting in brief openings. Increased membrane tension promotes upward movement of the TM4s due to changes in the shape of the channel that are energetically favored under tension. This movement of TM4 seals the lateral fenestrations, preventing lipid block, and results in longer duration and higher conductance mechanically-gated openings. Similar TM4 up and down states have been observed in the related mechanosensitive K2Ps TREK-1 and TREK-2, though gating in these channels also involves conformational changes in the K^+^-coordinating selectivity filter.

Given the physiological contribution of TRAAK and TREK-1 to action potential propagation, implication of TRAAK in epileptic disorders, and demonstrated channel inhibition by hydrophobic lipid acyl chains, we asked whether CBD modulates TRAAK—and related K2P channel—activity. We show that cannabinoids are potent blockers of mechanosensitive K2Ps and elucidate the mechanistic basis of channel inhibition.

## Results

We first investigated cannabinoid inhibition of TRAAK channels in excised patches. CBD potently inhibited TRAAK, with effects evident at 1-10 nM, full inhibition at ~1 µM, and an IC_50_ ~ 145 nM (Fig. 1A-D). Preliminary experiments using plastic perfusion components resulted in lower apparent potency, similar to effects reported in a study of CBD activation of K_v_7.2/7.3 channels and attributed to cannabinoid absorption to plastics ^16^. Final experiments were therefore performed with all glass components. We next asked if the related mechanosensitive K2Ps TREK-1 and TREK-2 were similarly cannabinoid-sensitive. All three channels were inhibited by CBD, but TREK-1 and TREK-2 were ~10-fold less sensitive than TRAAK (IC_50_s 1.5 and 1.3 µM, respectively, Fig. 1D-H). TREK-2 showed a steeper response profile and larger Hill-coefficient (1.3, 1.5, and 3.8 for TRAAK, TREK-1, and TREK-2, respectively), which suggests TREK-2 may have a unique CBD binding mode and/or stoichiometry.

**Figure 1 –.**
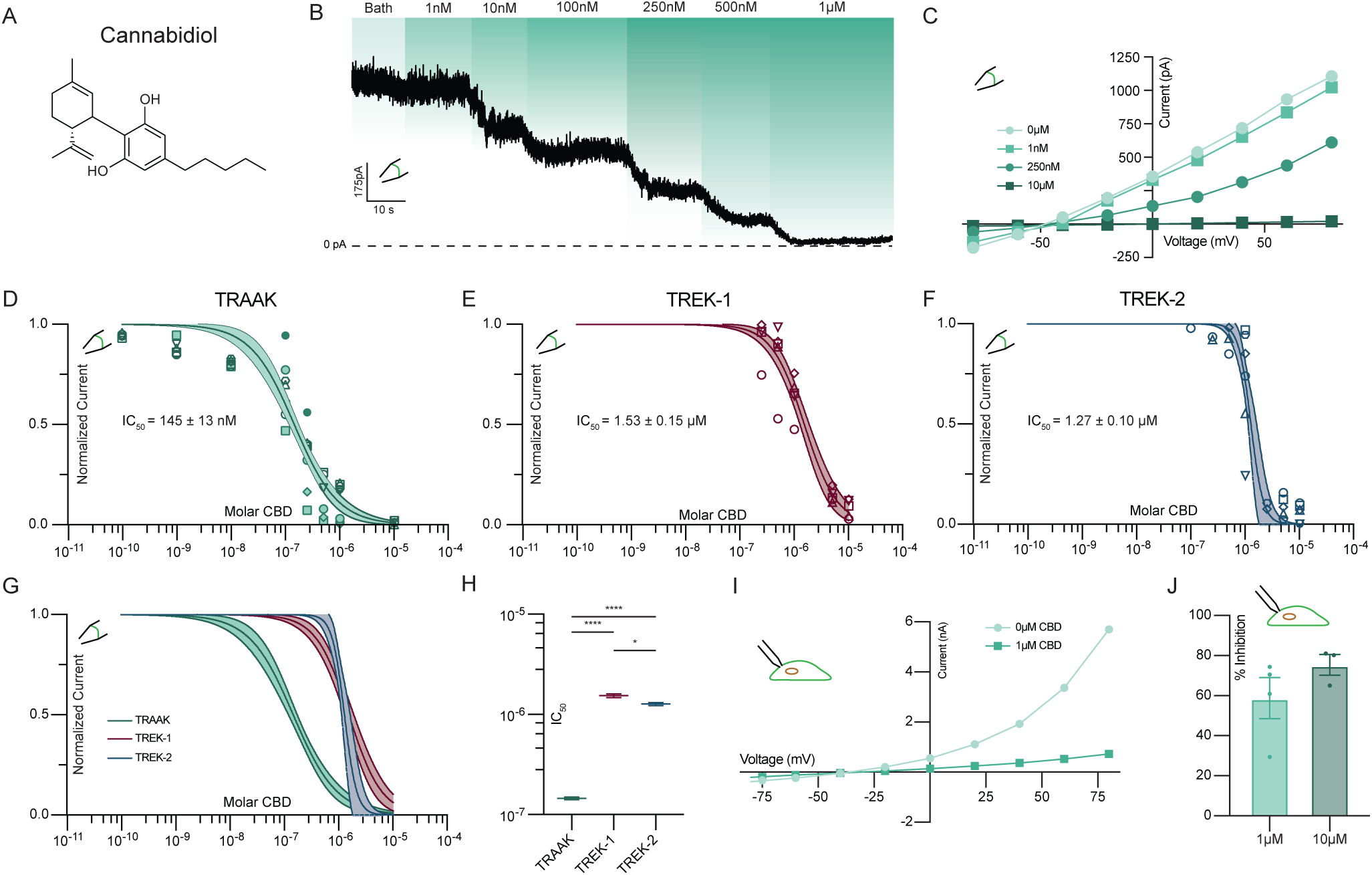
CBD inhibits mechanosensitive K^+^ channels. **(a)** Structure of cannabidiol (CBD). **(b)** Representative excised patch recording from a TRAAK-expressing cell perfused with increasing concentrations of CBD. Currents are recorded at 0 mV in a ten-fold K^+^ gradient. **(c)** Representative current-voltage relationship from TRAAK containing excised patch treated with increasing concentrations of CBD. **(d)** TRAAK, **(e)** TREK-1, **(f)** TREK-2, and **(g)** overlaid dose-response curves. Data from individual patches are plotted with different shapes. Solid lines are fits to four parameter logistic curves with 95% confidence intervals shaded. **(h)** Comparison of fit IC_50_ values (TRAAK: 145±4 nM, TREK-1: 1.53±0.06 µM, and TREK-2: 1.27±0.04 µM (mean ± sem, n = 9, 6, and 5 patches, respectively)). Differences assessed with Brown-Forsythe and Welch Annova with Dunnet correction for multiple comparisons. ****p<0.0001, *p=0.0295, n.s. not significant. **(i)** Representative current-voltage relationship and **(j)** average percent inhibition at 0mV from whole-cell recordings of TRAAK in response to 1 and 10 µM CBD.

In excised inside-out patch experiments, CBD has direct access to the intracellular side of the channels (Fig. 1A-H). To test whether CBD can inhibit TRAAK from extracellular side, we conducted whole-cell recordings from cultured cells. TRAAK remained strongly inhibited by extracellular CBD (Fig. 1I,J). While conducting these experiments, we found bath perfusion readily activated TRAAK, likely due to shear forces that increase membrane tension ^33,34^. We used focal perfusion and low solution flow rates to minimize mechanical channel activation, but cannot exclude residual mechanical effects contributing to the lower apparent sensitivity of TRAAK to extracellular compared to intracellular CBD (Fig. 1D,J).

CBD consists of a bicyclic head group and hydrocarbon tail (Fig. 1A). To identify the chemical features of CBD important for channel inhibition, we evaluated several related phytocannabinoids with select structural differences. Cannabinol (CBN), an oxidized derivative of Δ9-THC, has with the same hydrocarbon tail as CBD but a different, tricyclic head structure (Figure 2A). The additional aromatization in CBN increases head planarity and eliminates rotational freedom of motion relative to CBD. CBN is ~33% more potent than CBD (IC_50_ 107 nM), suggesting that the entropic cost of ordering the bicyclic head of CBD limits affinity. Cannabidiphorol (CBDP) and cannabidivarin (CBDV) share the same bicyclic head structure as CBD, but possess hydrocarbon tails with two additional or two fewer methylene units, respectively. These changes in hydrocarbon tail length have dramatic effects on potency. Compared to CBD, the more hydrophobic CBDP is ~10-fold more potent (IC_50_ 17±3 nM) while the less hydrophobic CBDV is ~40-fold less potent (5.58 ± 0.34 µM, Fig. 2B-D). CBDV still fully inhibits TRAAK channels at high (20 µM) concentration. This demonstrates that increasing planarity or hydrophobicity among related cannabinoids increases inhibitor potency.

**Figure 2 –.**
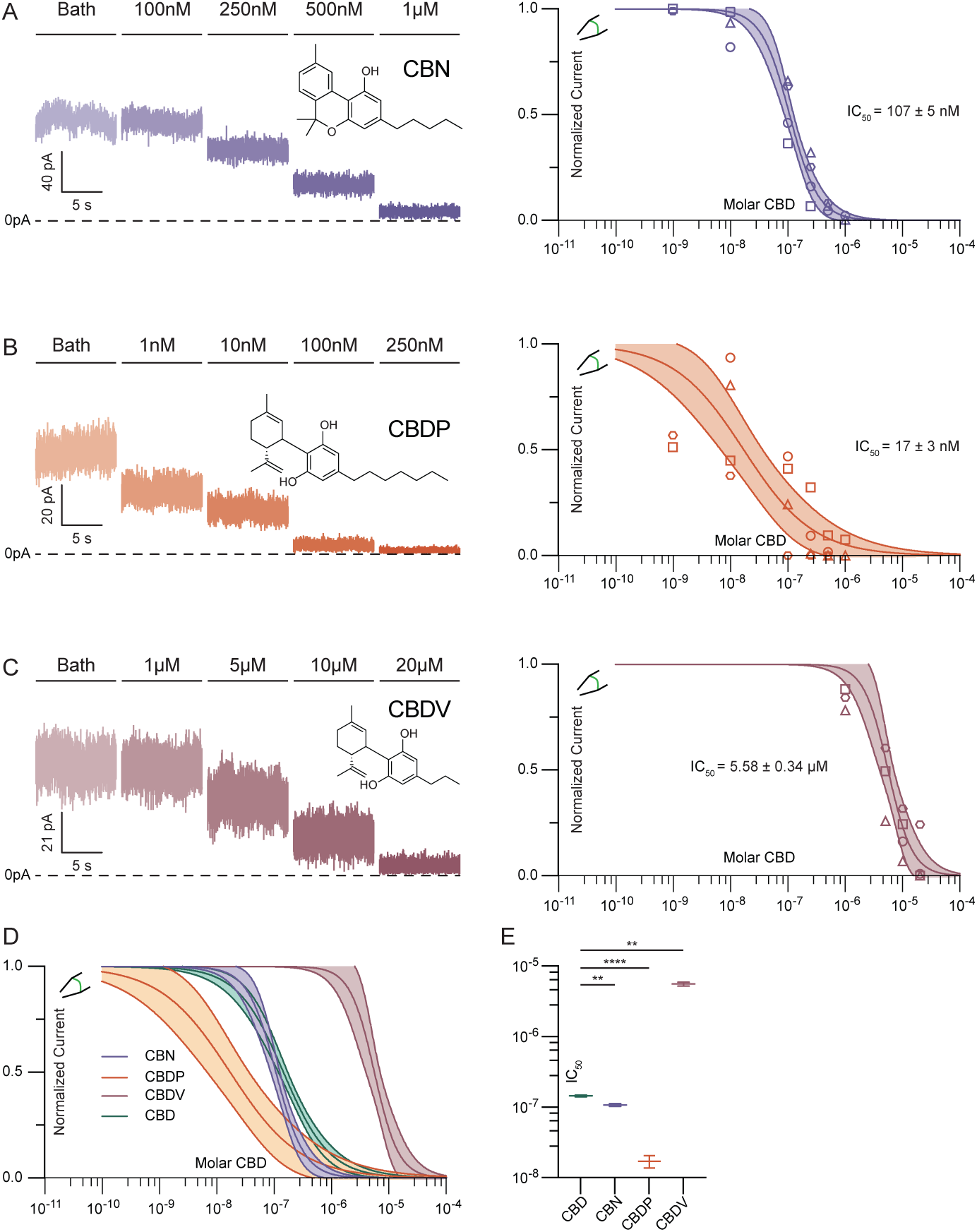
Potency of TRAAK inhibition is correlated with cannabinoid hydrophobicity. Representative recording (left) and dose response curve (right) from an excised patch of TRAAK treated with increasing concentrations of **(a)** CBN, **(b)** CBDP, and **(c)** CBDV. Data from individual patches are plotted with different shapes. Temporally discontinuous portions of the recording are indicated with gaps separating current trace. **(d)** Overlaid dose-response curves and **(e)** comparison of fit IC_50_ values (CBD: 145±4 nM, CBN: 107±5 nM, CBDP: 17±3 nM, and CBDV: 5.58 ± 0.34 µM (mean ± sem, n = 9, 4, 4, and 4 patches, respectively)). Differences assessed with Brown-Forsythe and Welch Annova with Dunnet correction for multiple comparisons. **p<0.01, ****p<0.0001, n.s. not significant.

Gain-of-function TRAAK mutations have been identified in human disease ^23–28^ and functional studies of K2Ps ^35,36^. The mechanistic basis for channel activation by several of these mutations is known and involves stabilization of either leak- or mechanically gated-like open states ^32^. We evaluated five TRAAK gain-of-function mutants to better understand CBD’s mechanism of action and to explore the susceptibility of disease-causing channel variants to cannabinoid inhibition.

We first considered the FHEIG-inducing mutations TRAAK_A270P_ and TRAAK_A198E_ ^23–25,28^. These mutants are highly activated under low-tension basal conditions, with open probabilities well over 0.9, because they mimic the effect of high membrane tension on wild-type channels ^32^. Both mutations shift the conformational equilibrium from the TM4-down, lipid-blocked/leak open state to TM4-up, mechanically gated-like open conformations ^32^. We reasoned that if CBD inhibits TRAAK by competing with lipid acyl chains to block the channel cavity in the TM4-down state, these mutants will show diminished sensitivity to inhibition. Indeed, TRAAK_A270P_ and TRAAK_A198E_ were ~8- and ~4-fold less sensitive than wild-type TRAAK to CBD (IC_50_ = 1.12±0.04 µM and 496±26 nM, respectively, Fig. 3A,B, Fig. S1). For both mutants, high (10 µM) concentrations of CBD fully inhibited channel activity.

**Figure 3 –.**
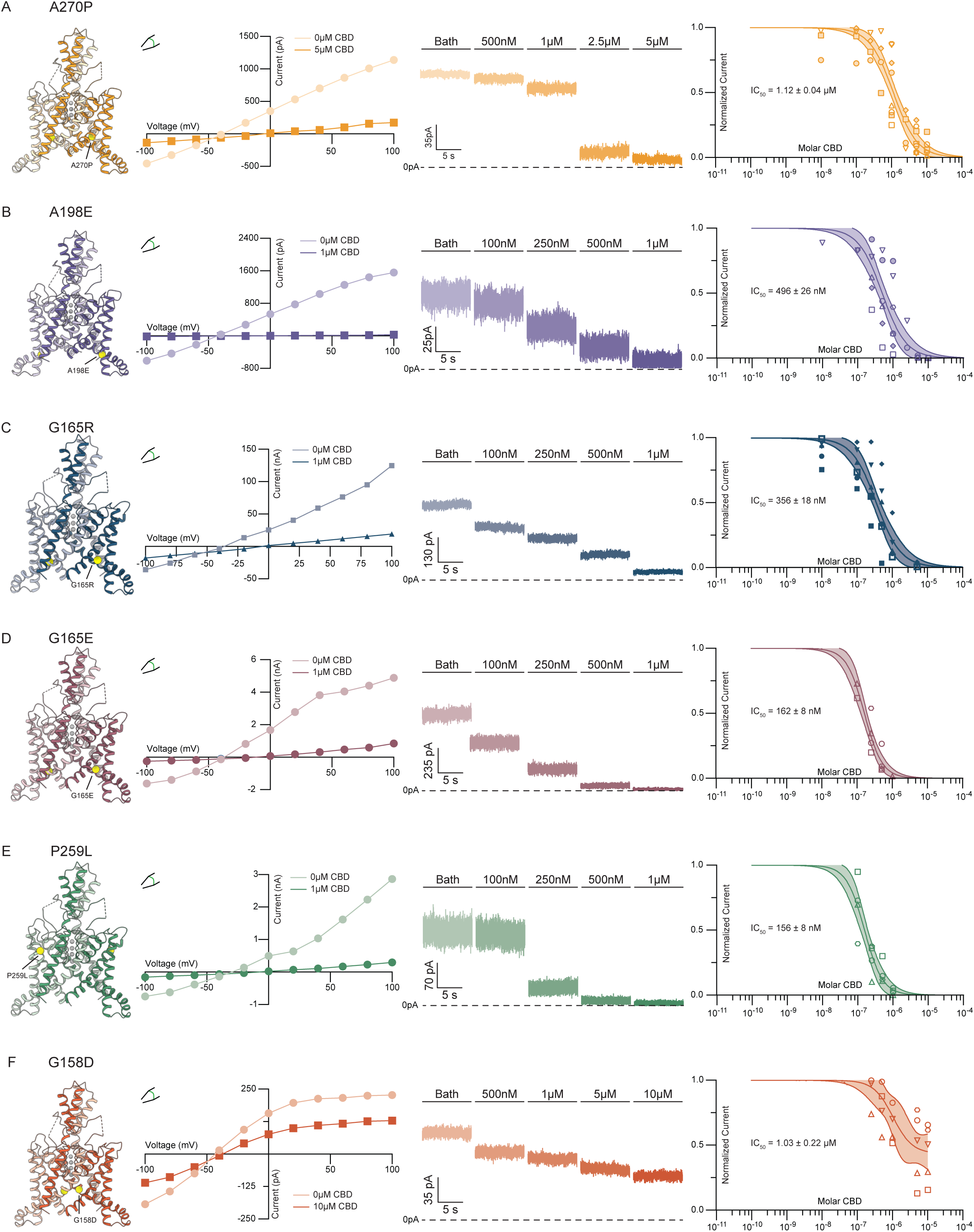
CBD inhibits disease-causing TRAAK variants. Inhibition of (a) TRAAK_A270P_, (b) TRAAK_A198E_, (c) TRAAK_G165R_, (d) TRAAK_G165E_, (e) TRAAK_P259L_, and (f) TRAAK_G158D_ by CBD. Determined or modeled structure viewed from the membrane plane with mutated residue highlighted in yellow, representative current-voltage relationship and current recording from an excised patch, and dose-response curve are shown from left to right. Data from individual patches are plotted with different shapes.

We next considered three mutants, TRAAK_G165R_, TRAAK_G165E_, and TRAAK_P259L_, recently identified in cases of FHEIG (G165R^26^, G165E^27^) and Rolandic epilepsy (P259L^27^). All three mutants retained high sensitivity to CBD (Figure 3C-E), with TRAAK_G165E_ and TRAAK_P259L_ statistically indistinguishable (IC_50_ = 162 ± 8 and 156 ± 8 nM, respectively) and TRAAK_G165R_ only modestly reduced compared to wild-type (IC_50_ = 356±18 nM) (Fig. 3C-E, Fig. S1). This suggests these mutations activate TRAAK to a lesser degree or in a manner distinct from the FHEIG-causing mutations TRAAK_A270P_, TRAAK_A198E_.

Finally, we considered TRAAK_G158D_. TRAAK_G158D_ is activated under basal conditions with an open probability ~0.7 and similar mutations at this site activate all vertebrate K2Ps ^32,35^. G158 points into the channel cavity towards the lipid binding site; introduction of the negatively charged aspartic acid electrostatically disfavors lipid block, shifting the conformational equilibrium towards a TM4-down, leak-like open state under basal conditions ^32^. We reasoned that if CBD, like lipids, binds in the channel cavity to block conduction, this mutant would be less sensitive to inhibition. The response of TRAAK_G158D_ to CBD treatment was found to be different from wild-type TRAAK in two ways. First, the channel was ~7.1-fold less sensitive to CBD with an EC_50_ = 1.03 ± 0.22 µM (Fig. 3F, Fig. S1). Second, CBD showed lower efficacy against TRAAK_G158D_ than wild-type TRAAK or other disease-causing variants, with inhibition plateauing at ~45% of initial current even at the highest CBD concentration tested (10 µM, Fig. 3E). These results suggest that (i) CBD has reduced affinity for TRAAK_G158D_ due to electrosteric repulsion between the mutated residue and the cannabinoid and (ii) CBD only partially occludes its conduction pathway, potentially because of an altered CBD binding pose.

Data to this point are consistent with a model in which cannabinoids bind in the channel cavity to inhibit mechanosensitive K^+^ channels and preferentially bind TM4-down over TM4-up states. A prediction of this model is that increasing membrane tension will reduce CBD inhibition of TRAAK current by promoting TM4-up states. We tested this prediction by using methyl-beta-cyclodextrin (mβCD) to increase membrane tension in TRAAK-containing patches. mβCD has been shown to irreversibly increase tension and activate force-gated ion channels by sequestering cholesterol and lipids from the membrane ^37,38^. Fig. 4A shows an experiment in which TRAAK activity and CBD sensitivity were recorded before and after treatment with mβCD. Prior to mβCD treatment, 1 µM CBD treatment resulted in near complete (91 ± 3%) inhibition (Fig. 4A,B) that was reversible after washout. Subsequent mβCD treatment activated channels and activity remained elevated following mβCD washout, indicating stable high membrane tension in the patch. From this activated state, 1 µM CBD treatment resulted in only partial (43 ± 13 %) inhibition. Approximately 10-fold higher CBD concentrations were required to achieve a similar degree of inhibition to prior to mβCD treatment (Fig. 4A).

**Figure 4 –.**
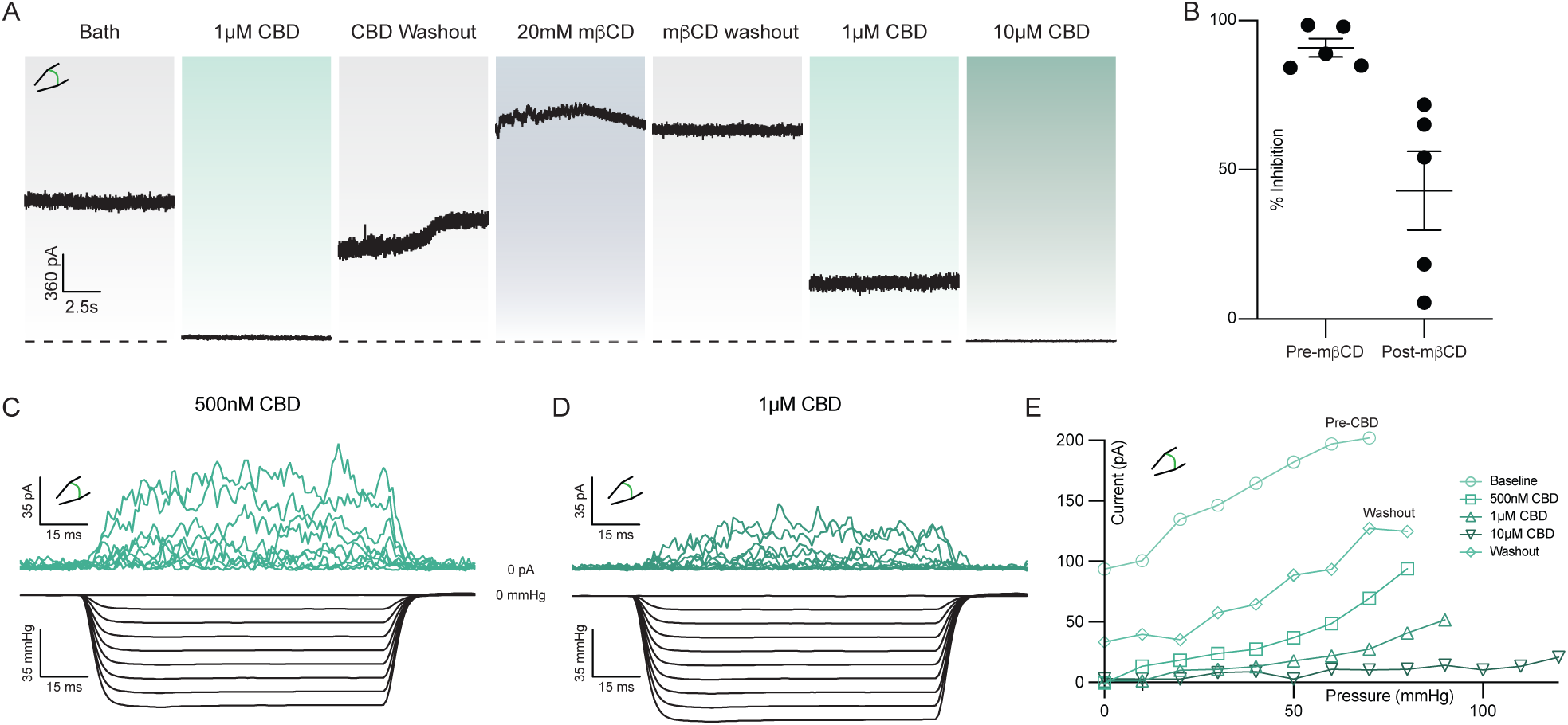
Mechanical activation of TRAAK reduces CBD efficacy. **(a)** Representative excised patch recording of TRAAK in response to CBD before and after treatment with methyl-β-cyclodextrin (mβCD) to chemically increase membrane tension. **(b)** Comparison of percent inhibition by 1 µM CBD before and after treatment with mβCD. Percent inhibition was calculated relative to the current prior to delivery of each dose of 1µM CBD. **(c,d)** Currents (top) recorded in response to steps of negative pressure (bottom) at **(c)** 500nM and **(d)** 1µM CBD. **(e)** Current versus pressure relationships from a representative recording before CBD treatment, during treatment with increasing concentrations of CBD, and after washout.

A second prediction of our model is that CBD inhibition will reduce mechanically activated TRAAK currents by preferentially binding to and stabilizing TM4-down conformations. To test this, we measured current elicited by steps of negative pressure before CBD application and after addition of 0.5, 1, and 10 µM CBD (Fig. 4C-E, Fig. S2). We found that as CBD concentration increased, more negative pressure was required to activate TRAAK to the same degree.

These results support our model for cavity binding and steric occlusion of the conduction path by CBD, but do not fully exclude an alternative explanation in which CBD inhibits TRAAK indirectly by decreasing basal tension or changing other membrane properties to promote closed channel states. We therefore asked if increasing concentrations of CBD alters membrane tension. In a patch recording, the Young-Laplace equation T = ΔPr/2 relates membrane tension (T) to the measurable pressure difference across the lipid bilayer (ΔP) and membrane radius of curvature (r). We measured changes in patch radius elicited by pressure steps before and after CBD treatment (Figure S3). We found that even 50 µM CBD, a concentration higher than that used in the other electrophysiology experiments herein, had no effect on membrane tension, consistent with direct inhibition of TRAAK and TREK channels by CBD.

We next pursued a structural approach to gain molecular insight into the basis for cannabinoid inhibition of mechanosensitive K^+^ channels. Cryo-EM structures of *Danio rerio* TREK-1 in apo, phosphatidylethanolamine(PE)-inhibited, and phosphatidic acid (PA)-activated states are known ^39^. Like human TREK-1 (Fig. 1E), we found *Danio rerio* TREK-1 is inhibited by CBD (IC_50_ = 11.1 ± 1.3 µM, Supplementary Fig. 4). We determined a cryo-EM structure of *Danio rerio* TREK-1 in the presence of 100 µM CBD (Fig. 5A-B, Figs. S5–6, Table S1). The structure was resolved to an overall resolution of 3.94 Å, though local resolution for most of the channel is substantially higher and map quality in regions discussed below is comparable to previously reported structures resolved to 2.8-3.5 Å ^39^ (Fig. S6).

**Figure 5 –.**
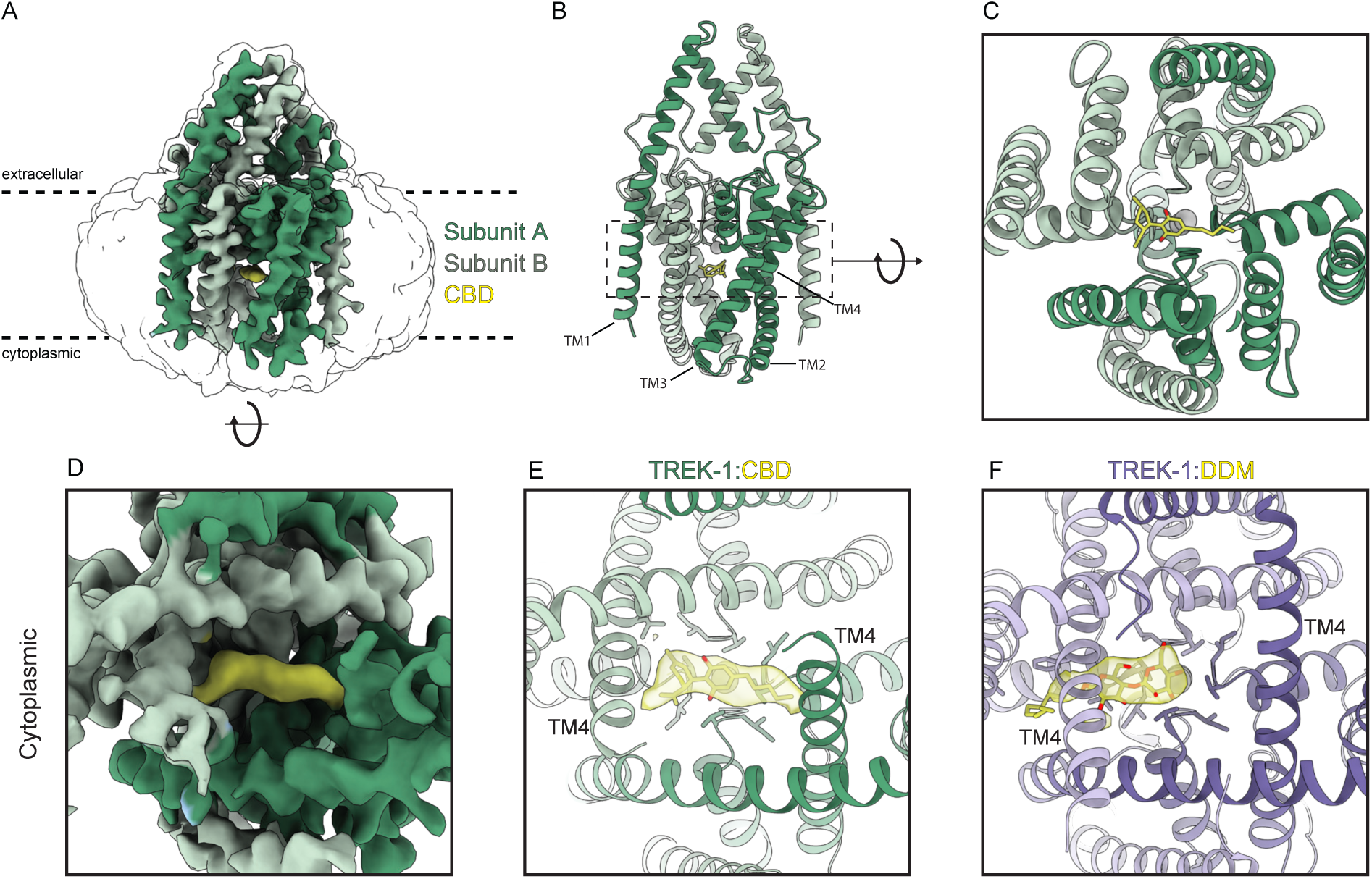
Cryo-EM structure of TREK-1 bound to CBD. **(a)** 3.9 Å resolution cryo-EM map of *Danio rerio* TREK-1 in complex with CBD viewed from the membrane. TREK1-subunits are green and CBD is yellow. Unsharpened map is shown transparent at low contour. Overall model is shown from **(b)** the membrane plane and **(c)** the cytoplasmic side. **(d)** Zoomed view of sharpened map from the cytoplasmic side. **(e,f)** Comparison of models with cavity-occupying molecule and cryo-EM density for **(e)** TREK-1:CBD and **(f)** apo TREK-1 structure. CBD and dodecyl maltoside detergent occupy the cavity in the current and previously reported apo structures, respectively.

A strong, elongated density feature consistent with CBD is evident in the channel cavity directly underneath the selectivity filter (Fig. 5A-E). The density is best fit with the CBD aromatic ring centered under the filter and the terpene head group and hydrophobic tail projecting on either side towards the lateral fenestrations. We note this placement is speculative because the resolution is insufficient to unambiguously define CBD binding pose. Several lines of structural evidence support this model for CBD binding in the channel cavity. First, cavity density in the TREK-1:CBD structure is markedly different than that in the apo TREK-1 structure ^39^, which was prepared in the same way apart from the addition of CBD. In apo TREK-1, the cavity density is smaller and biased towards one subunit, extends into the surrounding micelle, and is consistent with a modeled dodecyl maltoside detergent (DDM) molecule (Fig. 5E,F). Second, CBD addition resulted in conformational changes to TM4 position. In the TREK-1:CBD structure both TM4s are down, while in apo TREK-1 one TM4 is up and one TM4 is down (Fig. 5E, Fig. S7). The symmetric TM4-down TREK-1:CBD conformation is more similar to the inhibited TREK-1:PE structure than the apo TREK-1 structure (Fig. S7). Third, the TREK-1:CBD structure shows differences in selectivity filter occupancy (Supplementary Fig. 8). In the presence of CBD, density for K^+^ is evident in sites 2 and 4. In contrast, K^+^ density is evident in sites 1-3 in apo TREK-1, 1-3 in TREK-1:POPE, and 1-4 in TREK-1:POPA structures ^39^. We conclude CBD inhibits TREK-1 by binding in the channel cavity to sterically block ion passage. Destabilization of K^+^ coordination in the selectivity filter may additionally contribute to channel inhibition.

## Discussion

Data presented here support a model in which CBD inhibits mechanosensitive K^+^ channels by binding in the channel cavity to block ion conduction (Fig. 6). At low tension, TRAAK and TREK channels predominantly adopt TM4-down conformations that expose lateral fenestrations above TM4 to the surrounding lipid membrane ^31,32^. In this conformation, the channels are predominantly closed because lipid acyl chains access the cavity through lateral fenestrations and block ion conduction; spontaneous delipidation results in low leak activity. High tension promotes a TM4-up conformation that closes lateral fenestrations to prevent lipid block, resulting in a high conductance, open channel. We found increasing mechanical activation of TRAAK reduces CBD inhibition, while increasing CBD concentration reduces mechanically activated currents (Fig. 4). Mutations that promote TM4-up conformations decrease channel sensitivity to CBD (Fig. 3). The TREK-1:CBD structure shows both TM4s down, rather than one up and one down as in the apo TREK-1 structure ^39^. This is likely because CBD as modeled would sterically clash with TM4 residues if the helix adopted an up conformation. We conclude CBD preferentially binds to TM4-down channel conformations and effectively competes with abundant membrane lipids for an overlapping cavity site to block ion conduction. Whether CBD accesses the channel cavity through membrane-facing fenestrations, through the cytoplasm, or both, remains to be determined.

**Figure 6 –.**
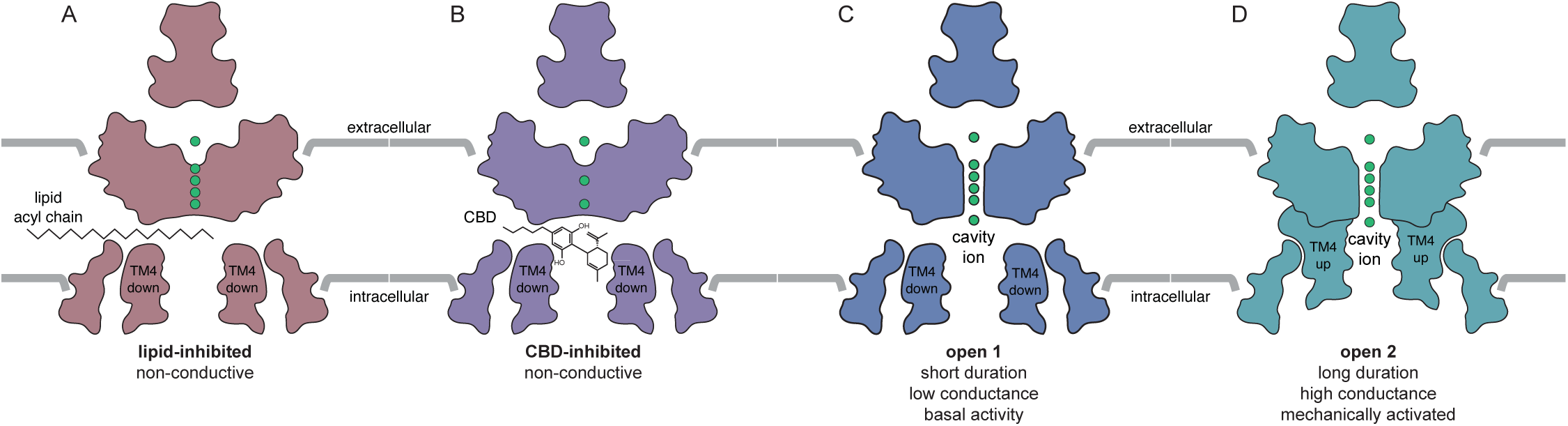
Model for CBD inhibition of mechanosensitive K^+^ channels. CBD binds in the channel cavity to sterically occlude ion conduction. CBD preferentially binds to TM4-down states (lipid-blocked and leak open states) over the TM4-up mechanically-gated open state ^32,36,39^.

CBD modulates distantly-related ion channels in the voltage-gated ion channel superfamily, inhibiting Na_v_s and BK ^13,18^ and activating TRPVs, TRPA1 ^14,15^, and KCNQs ^16^. With an IC_50_ of 145 nM, TRAAK is among the highest-affinity CBD targets known, while TREK-1 and TREK-2 show roughly 10-fold lower potency similar to Na_v_s. CBD is the highest affinity TRAAK inhibitor reported to date with ~3-5 fold higher affinity than RU-TRAAK-1 and -2 and greater than 10-fold higher affinity than nonspecific inhibitors ruthenium red and TKDC^40,41^.

Structures of TRPV2 ^15,42,43^, Na_v_1.7 ^44^, and KCNQ7.2/7.3 ^17^ in complex with CBD have been reported. Notably, CBD binds all these channels in similar fenestration site(s) that connect channel cavities to the surrounding membrane. CBD binding within all four fenestrations of TRPV2 and KCNQ channels stabilizes their open states, while binding within one fenestration of Na_v_1.7 (together with a second CBD molecule binding to the IFM motif) stabilizes the inactivated state. In bacterial Na_v_M, CBD binds deeper within all four fenestrations, partially entering the channel cavity ^45^. The mechanism of action of CBD on mechanosensitive K^+^ channels is fundamentally different. CBD does not bind within fenestrations, but rather fully accesses the channel cavity to bind directly underneath the selectivity filter (Fig. 5). Differences in these binding sites suggests structural insights could guide the design of cannabinoid derivatives with higher specificity to particular ion channels.

CBD inhibition of mechanosensitive K2Ps has potential clinical relevance that warrants further exploration. The molecular target of CBD that underlies its antiepileptic effects remains unknown, but TRAAK is a compelling candidate for several reasons. TRAAK activity controls spike propagation in myelinated neurons, where it localizes to axon initial segments ^19,20^ and nodes of Ranvier ^21,22^. In these specialized neuronal compartments, TRAAK colocalizes with Na_v_ channels, the target of most classic anticonvulsants, and KCNQ channels, the target of the only approved K^+^ channel-specific antiepileptic. Inhibition of TRAAK or TREK-1 reduces Na_v_ availability by preventing recovery from inactivation, thereby limiting spike velocity and frequency^21,22^, which could diminish the ectopic excitability characteristic of epileptic disorders. In addition, gain-of-function mutations in TRAAK cause epilepsy in humans, either in a form of Rolandic epilepsy or the neurodevelopmental disorder FHEIG ^23–28^. While some of these patients respond to anti-seizure medications including carbamazepine, oxcarbazepine, and valproate, others are refractory to treatment ^26^. There are currently no specific TRAAK inhibitors known. We show all known disease-causing variants (TRAAK_G165E_, TRAAK_G165R_, TRAAK_P259L_, TRAAK_A270P_, and TRAAK_A198E_) retain sensitivity to inhibition by CBD. These results suggest CBD or other cannabinoids could be pursued as a targeted treatment of TRAAK channelopathies in patients.

## Methods

### Electrophysiology

For patch clamp recording from *Xenopus* oocytes, genes encoding full-length *Homo sapiens* TRAAK (UniProt Q9NYG8-2), TREK-1 (UniProt O95069), and TREK-2 (UniProt P57789) were codon optimized for eukaryotic expression, synthesized (Genewiz), and cloned into a modified pGEMHE vector using Xho1 and EcoR1 restriction sites. The transcribed messages encode *H. sapiens* TRAAK amino acids 1–393, TREK-1 amino acids 1–426, or TREK-2 amino acids 1–538 with an additional three amino acids (SNS) at the C terminus. TRAAK mutants were introduced by polymerase chain reaction (PCR). Linearized DNA was transcribed *in vitro* using T7 polymerase. Complementary RNA (0.1 to 10 ng for TRAAK, TREK-1, TREK-2, and mutants) in 50 nL H2O was injected into *Xenopus laevis* oocytes extracted from anesthetized frogs. Currents were recorded at 25 °C from inside-out patches excised from oocytes 1 to 5 d after RNA injection. The pipette solution contained 15 mM KCl, 135 mM NaCl, 2 mM MgCl_2_, and 10 mM HEPES (pH = 7.4 with KOH) and the bath solution contained 150 mM KCl and 10 mM HEPES (pH = 7.1 with KOH). Borosilicate glass pipettes were pulled to 2-5 MΩ resistance.

For patch clamp recording from cultured cells, *dr*TREK-1 or *hs*TRAAK genes were cloned into a modified pCEH vector to generate C-terminally EGFP-tagged constructs^29^. HEK293T were cultured in DMEM (Gibco, Thermo Fisher Scientific) with 10% FBS and 100 U ml−1 penicillin and 100 μg ml−1 streptomycin. Trypsinized cells were deposited on 12-mm glass coverslips in a six-well dish one to two days before transfection. For transfection, *dr*TREK-1 or *hs*TRAAK plasmids were mixed with FuGENE 6 in OptiMEM at a 1:3 ratio with 1 μg DNA per well per construct and applied to cells in growth medium with antibiotics. Patching was conducted from 24 to 60 h post-transfection. Coverslips were placed in a perfusion chamber at room temperature in isotonic bath solution: 135 mM NaCl, 15 mM KCl, 1 mM CaCl_2_, 1 mM MgCl_2_, 10 mM HEPES, adjusted to a final pH of 7.5 with NaOH. Cells were chosen by a combination of the presence GFP fluorescence and cell morphology consistent with healthy interphase cells. Borosilicate glass pipettes were pulled to a resistance of 1.2-1.8 MΩ for patch electrophysiology and 1.8-2.8 MΩ for whole cell electrophysiology. Pipettes were filled with pipette solution: 150 mM KCL, 5 mM EGTA, 3 mM MgCl_2_, and 10 mM HEPES, adjusted to pH 7.5 with KOH.

Patches were evaluated with voltage-step protocols to confirm TRAAK expression and minimal non-specific leak based on reversal potential. Currents were recorded using an Axopatch 200B amplifier (Molecular Devices) at a bandwidth of 1 kHz and digitized with an Digidata 1550B (Molecular Devices) at 100-500 kHz using pClamp10.7 software. Pressure was applied with a second-generation high-speed pressure clamp device (HSPC-2-SB, ALA Scientific Instruments). Aggregated data were from patches from n ≥ 2 different cells on different days. No relevant differences were observed between patches from different cells. In an effort to minimize variability associated with differences in resting membrane tension, we targeted patches with low basal curvature and low resting channel activity and ensured measured pressure difference across the patch was as close to zero as possible at rest to prevent mechanical activation of channels. Data were analyzed using Clampfit 10.7, Excel, and Graphpad Prism software. For voltage families, the data were decimated 100x and plotted in Prism. Statistical analysis was conducted with Dunnett’s multiple comparisons test after an ordinary one-way ANOVA using Prism software.

To avoid cannabinoid absorption to plastic, cannabinoid solutions were made in glass vials and diluted by weight, perfusion was performed using glass tubing pulled from borosilicate glass Pasteur pipettes (Fisherbrand, Cat# 13-678-20C), and recordings were performed in glass chambers. Channel activity was continuously recorded during perfusion steps. Patches were exposed to each cannabinoid concentration until a stable current level was reached (0.5-2 minutes). CBD (Cayman Chemical cat#90080) was prepared as a 10mM stock solution in DMSO, CBN (Cayman Chemical cat#25495) was prepared as a 80mM stock solution in DMSO, CBDV (Phytolab Cat#85955) was prepared as a 10mM stock solution in DMSO, and CBDP (Cayman Chemical cat#33611) was purchased as a 2.9 mM stock solution in methanol. Cannabinoids were serially diluted in bath solution to achieve working concentrations of 50 µM to 10 nM. Bath volume in the recording chamber was minimized by suction and drug-containing solutions were delivered three times to ensure establishment of desired concentration in the recording chamber. Cannabinoids were applied from low to high concentration. The recording chamber and perfusion system were washed extensively with methanol after every trial.

The effects of cannabinoids on channel activity were quantified by the reduction in channel current evoked at 0 mV during a gap-free recording or during 200 ms long voltage families with steps ranging from −100 to +100 mV. Currents were normalized to those prior to cannabinoid application to facilitate comparisons across different patches and cells. Dose response curves were four-parametric Boltzmann fits to aggregated data. For voltage-step recordings, current was measured at 0mV from each recording. Data were aggregated for all recordings with and without drug present and were normalized to the baseline activity before drug administration. Comparisons between constructs and drug isoforms were conducted with ordinary one-way ANOVA and Tukey multiple-comparison tests.

### Patch imaging and membrane tension calculation

A construct encoding EGFP fused to the CAAX-containing C-terminal tail of H. sapiens H-Ras (NP_005334 amino acids 170–189) through a GGRS linker was cloned into a pCS2+ vector using Gibson assembly. Linearized DNA was transcribed *in vitro* using T7 polymerase. 3– 10 ng complementary RNA in 50 nL H2O was injected into *Xenopus laevis* oocytes. Excised patches were illuminated with an LED light engine (SpectraX, Lumencor) through a GFP filter (450/50 nm excitation, 506 nm dichroic mirror, 500 nm longpass emission filter) and water immersion objective lens (x60, NA1.0). Movies were recorded at 120 Hz with an infrared camera (IR-2000, DAGE-MTI). Images were preprocessed within FIJI (ImageJ). Image contrast was enhanced to facilitate analysis. Video files were loaded into FIJI and converted into a JPEG stack. Frames were time matched to stimuli by multiplying frame rate and time. Tension was calculated using python scripts as previously reported ^30^.

### TREK-1 expression and purification

*Danio rerio* TREK-1 (UniProt Q9NYG8-2) was cloned for expression in *Pichia pastoris* as previously described ^29^ with modifications described here. The construct used for purification included an additional 26 amino acid N-terminal sequence from human TRAAK compared to Q9NYG8 that improved heterologous expression. The final construct is C-terminally truncated by 119 amino acids, incorporates two mutations to remove N-linked glycosylation sites (N104Q/N108Q), and is expressed as a C-terminal PreScission protease-cleavable EGFP-10x His fusion protein. As a result, there is an additional amino acid sequence of ‘‘SNSLEVLFQ’’ at the C terminus of the final purified protein after protease cleavage.

Starter cultures of recombinant Pichia were grown in YPD with 0.5 mg/mL Zeocin and grown at 30°C with shaking at 250 rpm overnight (12-14 h). Four 1L flasks of BMGY with 25ug/mL Zeocin were inoculated with 10mL of starter culture each. Cells grew for approximately 24 hours at 30°C at 250 rpm. 1L cell cultures were pelleted by centrifugation at 8000g for 10 minutes and resuspended in 1L BMMY with 25ug/mL Zeocin. Cells were harvested 40-60 h after induction with methanol and flash frozen in liquid N_2_. ~60 g of frozen *Pichia* cells expressing *dr*TREK-1 were disrupted by milling (Retsch model MM301) 5 times for 3 min at 25 Hz. All subsequent purification steps were carried out at 4°C. Milled Pichia cells were thawed in 200 mL of Lysis Buffer containing 50 mM TRIS, 150 mM KCl, 1mM EDTA pH 8. Protease inhibitors (Final Concentrations: E64 (1 μM), pepstatin A (1 μg/mL), soy trypsin inhibitor (10 μg/mL), benzamidine (1 mM), aprotinin (1 μg/mL), leupeptin (1μg/mL), AEBSF (1mM), PMSF (1mM)), benzonase (10µL) and DNAse (10 µL) were added to the lysis buffer immediately before use. Cells were lysed by sonication and centrifuged at 150,000 x g for 45 minutes. The supernatant was discarded, and residual nucleic acid was removed from the top of the membrane pellet using DPBS. Membrane pellets were scooped into a Dounce homogenizer containing extraction buffer (50 mM TRIS, 150 mM KCl, 1 mM EDTA, 1.5% n-Dodecyl-β-D-Maltopyranoside (DDM, Anatrace, Maumee, OH), 0.3% cholesteryl hemisuccinate Tris salt (CHS, Anatrace, Maumee, OH) pH 8). A stock solution of 10% DDM, 2% CHS was dissolved and clarified by bath sonication in 200 mM HEPES pH 8 prior to addition to buffer to the indicated final concentration. Membrane pellets were then homogenized in extraction buffer and this mixture (150 mL final volume) was gently stirred at 4°C for 2 hours. The extraction mixture was centrifuged at 33,000 x g for 45 minutes and the supernatant, containing solubilized membrane protein, was bound to 4 mL of Sepharose resin coupled to anti-GFP nanobody for 2 hours at 4°C. The resin was then collected in a column and washed with 10 mL of buffer 1 (20 mM TRIS, 150 mM KCl, 1 mM EDTA, 0.025% DDM, 0.005% CHS, pH 8), 40 mL of buffer 2 (20 mM TRIS, 500 mM KCl, 1 mM EDTA, 0.025% DDM, 0.005% CHS, pH 8), and 10 mL of buffer 1. The resin was then resuspended in 6 mL of buffer 1 with 0.5 mg of PPX protease and rocked gently in the capped column overnight (~ 12–14 h). Cleaved TREK-1 was then eluted with an additional 12 mL of wash buffer, spin concentrated to ~1 mL with Amicon Ultra spin concentrator 100 kDa cutoff (Millipore) and loaded onto a Superose S200 increase column (GE Healthcare, Chicago, IL) on an NGC system (Bio-Rad, Hercules, CA) equilibrated in an elution buffer (20 mM TRIS, 150 mM KCl, 1 mM EDTA, 0.025% DDM, pH 8). Peak fractions containing TREK-1 protein were then collected and spin concentrated prior to sample freezing.

### Cryo-electron microscopy

TREK-1 in DDM detergent was prepared at a final concentration of 3.1 (purification 1) or 4.4 (purification 2) mg/mL. CBD was spiked into the sample from a stock concentration of 10 mM in DMSO to a final concentration of 100 µM. The sample was incubated on ice for 30 minutes then clarified by a 10-minute 21,000 x g spin at 4°C prior to grid preparation. 3.4 μl of protein was applied to freshly glow discharged Holey Carbon, 300 mesh R 1.2/1.3 gold grids (C-flat, Electron Microscopy Sciences, USA (purification 1), Quantifoil, Großlöbichau, Germany (purification 2)) and plunge frozen in liquid ethane using a FEI Vitrobot Mark IV (ThermoFisher Scientific) set to 4°C, 100% humidity, 1 blot force, wait time of ~5 seconds, and 3 second blot time. Grids were clipped and stored in liquid nitrogen.

Both datasets were collected on a Titan Krios G3i electron micro-scope (Thermo Fisher) operated at 300 kV and equipped with a Gatan BioQuantum Imaging Filter with a slit width of 20 eV. Dose-fractionated images (~50 electrons per Å^2^ over 50 frames) were recorded on a K3 direct electron detector (Gatan) at a pixel size of 0.848 Å. 242 movies were collected in an 11 × 11 hole pattern with two targets per hole around a central hole position using image shift. Defocus was varied from −0.5 to −1.8 μm using SerialEM ^46^.

Motion correction was performed on 9,046 micrographs from collection 1 and 8,470 micrographs from collection 2 using patch motion correction in cryoSPARC v4.4.1 ^47^. Contrast transfer function (CTF) parameters were fit with patch CTF ^48^. For collection 1, template-free auto-picking of particles was performed with a blob picker with a minimum particle size of 120 Å and a maximum particle size of 200 Å on 5,481 movies CTF fit to 5.0Å or better, yielding an initial set of 2,315,705 particles. These were extracted at a 300-pixel box size for two-dimensional (2D) classification. Iterative rounds of 2D classification yielded a set of 45,022 particles that were used to train a Topaz model. Topaz-trained particle-picking yielded 1,176,701 particles which were manually curated through iterative rounds of 2D classing and multiple multi-class Ab-initios in which particle subsets that yielded the best 3D structure were chosen. Final ab-initio and non-uniform refinements (C1, 4 extra passes, 15Å initial resolution) yielded a 5.14 Å map from 261,227 particles.

For collection 2, the topaz model from collection 1 was utilized to conduct topaz-trained particle picking on 7,968 movies CTF fit to 7Å or better. Topaz picking and extraction a box size of 300Å yielded an initial set of 1,673,778 particles. These particles were curated through iterative rounds of 2D classification to a subset of 129,260 particles. These particles were then re-extracted from 3729 micrographs CTF fit to between 2.4Å and 7Å for a subset of 92,936 particles that were then used to train a new topaz model on the 3729 micrographs CTF fit between 2.4Å and 7Å. The second round of Topaz training, picking, and extraction yielded 750,776 particles. These classes were manually curated through a single round of 2D classification before a 3-class Ab-initio. The class with the clearest protein density was chosen yielding a subset of 316,175 particles with clear protein density. These particles were subjected to iterative rounds of 2D and 3D classification. Final ab-initio and non-uniform refinements (C1, 4 extra passes, 15Å initial resolution) yielded a 5.86 Å map from 228,680 particles.

The 261,227 particles from collection 1 and 228,680 particles from collection 2 were then re-extracted at a 300Å box size and joined into one particle set. Particles too close to the edges of micrographs were discarded, resulting in a joint subset of 484,500 particles. These particles were curated over 3 rounds of 3D classification to remove remaining junk. An ab-initio reconstruction of the remaining 160,072 particles was performed to provide an initial volume and a subsequent non-uniform refinement (C1, 4 extra passes, 15Å initial resolution) which resulted in a map with a 4.31 Å overall resolution. This map was post-processed in cryoSPARC v4.4.1 and used for reference-based motion correction. The resulting ‘shiny’ particles were subject to one round of 2D classification before one 3-class 3D classification resulting in 117,795 particles. These particles were used to generate a new ab initio model. We generated a mask to refine this map in Chimera (UCSF) from the previously reported *dr*TREK-1 structure ^39^. This mask was imported to cryoSPARC v4.4.1 and modified to a dilation radius of 7 pixels and a soft padding of 25 pixels. Local refinement (C1, 2 extra passes, 12Å initial resolution) of our volume with this mask resulted in a map with 4.15Å overall resolution. This map was utilized for a second round of reference-based motion correction. An ab initio and local refinement (C1, 2 extra passes, 12Å initial resolution) of the twice-polished 117,609 particles resulted in a final map of 3.94Å nominal resolution.

The final non-uniformed local refined and sharpened map from Cryosparc was used for modeling. An initial apo TREK-1 model (PDB: 8DE7) was rigid body fit to the density in Phenix ^49^. Model building was performed iteratively with manual adjustment in Coot ^50^, global real space refinement in Phenix ^51^, and geometry assessment in Molprobity^52^.

## Acknowledgements

This work was supported by the NIGMS GM145869 (S.G.B.) and NIH/NINDS NS125102 (B.S.). We thank D. Toso, P. Tobias, and R. Thakkar at the Cal-Cryo Electron Microscopy Facility for help with data collection. We thank B. Wainger (Harvard Medical School) for discussions and members of the Brohawn lab for feedback on the manuscript.

## Author Contributions

T.A.D., B.S., and S.G.B. conceived of the project. T.A.D., B.S., and R.D. performed electrophysiological experiments and analyzed data. T.A.D. and C.D. performed protein expression, protein purification, cryo-EM data collection, and cryo-EM data processing. T.A.D. and S.G.B. performed structure modeling and refinement. T.A.D. and S.G.B. wrote the manuscript. S.G.B. secured funding.

## Declaration of Interests

SGB and TAD are inventors on a patent application filed by the University of California, Berkeley related to research described here.

## Data and material availability

The TREK-1:CBD model is deposited in the PDB under 9DBR. Maps are deposited in the Electron Microscopy Data Bank (EMDB) under EMD-46725. Original micrograph videos and final particle stack are deposited in the Electron Microscopy Public Image Archive.

---

**Supplementary Figure 1 –.**
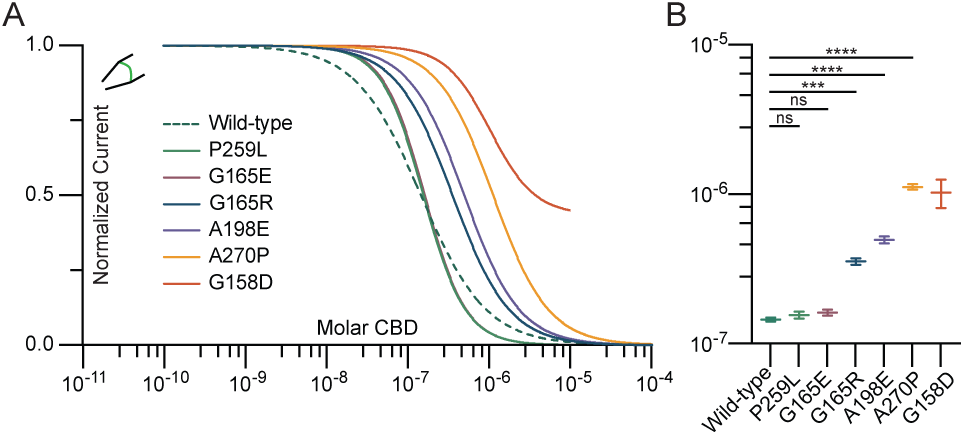
Comparison of CBD inhibition of disease-causing TRAAK variants. **(a)** Overlaid dose-response curves and **(b)** comparison of fit IC_50_ values (wild-type: 145±4 nM, P259L: 156±8 nM, G165E: 162±8 nM, G165R: 356±18 nM, A198E: 496±26 nM, A270P: 1.12±0.04 µM, and G158D: 1.03 ± 0.22 µM (mean ± sem, n = 9, 4, 4, 6, 7, 8 and 5 patches, respectively). Differences assessed with Brown-Forsythe and Welch Annova with Dunnet correction for multiple comparisons, ***P<0.001, ****p<0.0001, n.s. not significant.

**Supplementary Figure 2 –.**
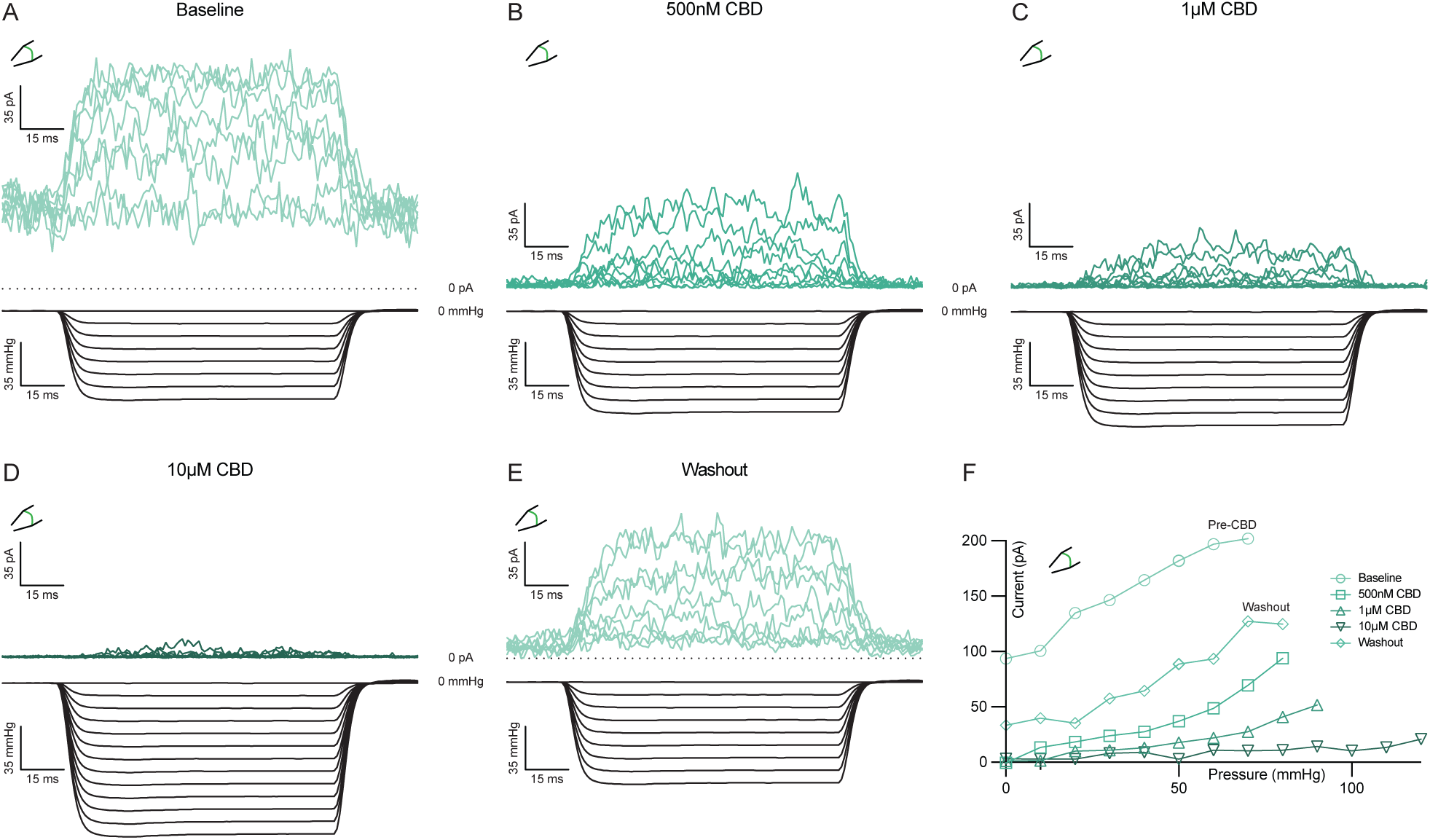
CBD inhibition reduces mechanical activation of TRAAK. **(a-e)** Example current traces from a TRAAK-containing excised patch in response to negative pressure at varying concentrations of CBD ranging from **(a)** 0µM, **(b)** 500nM, **(c)** 1µM, **(d)** 10µM, and **(e)** following washout. **(f)** Peak current versus pressure from data in **(a-e)**.

**Supplementary Figure 3 –.**
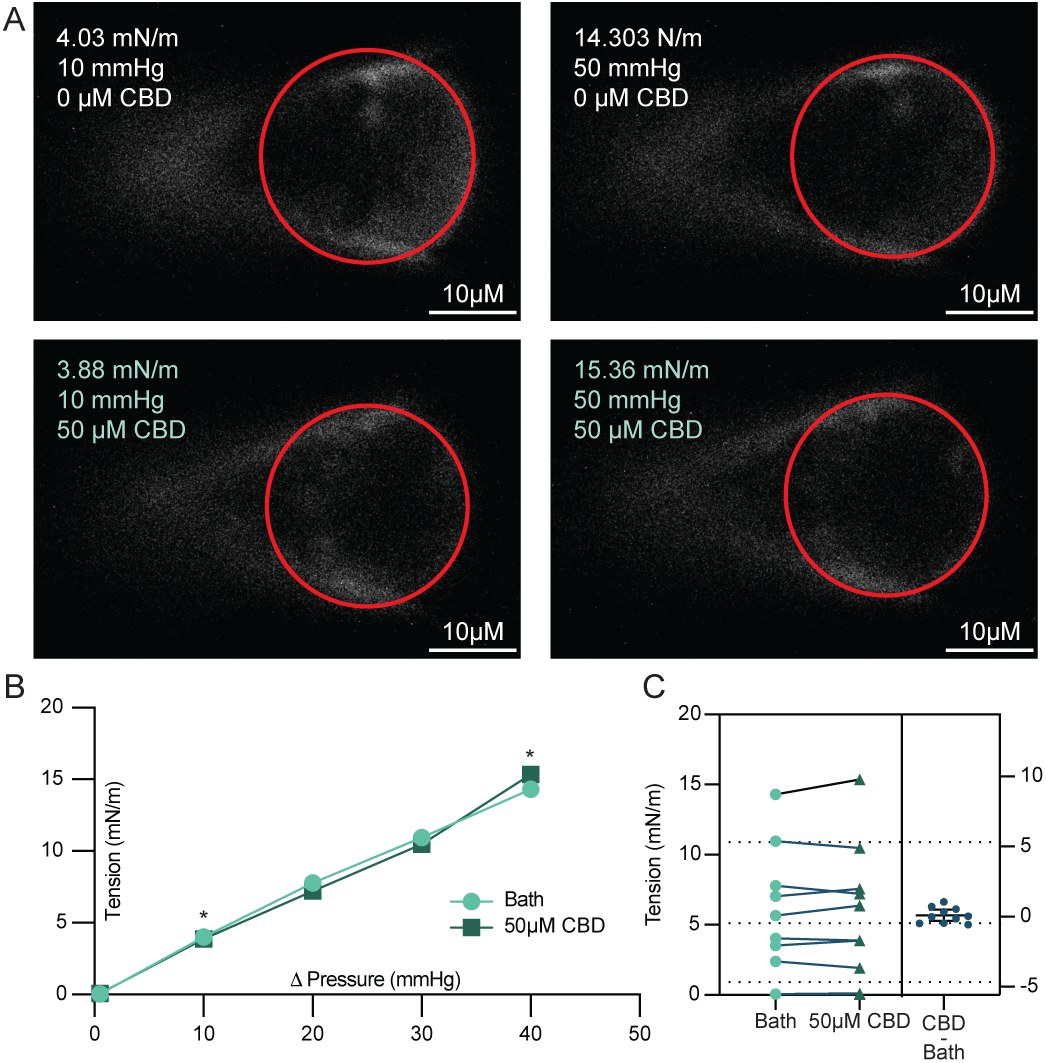
CBD treatment does not alter membrane tension. **(a)** Representative images of membrane patches under −10 (left) and −50 mmHg (right) pressure in 0 µM CBD (top) and 50µM CBD (bottom). **(b)** Calculated membrane tension versus applied negative pressure in 0 µM CBD (light green) and 50 µM CBD (dark green). Asterisks indicate data from images shown in **(a). (c)** Comparison of tension generated at matching pressure steps. Data from 10 paired recordings from three different patches are shown (p = 0.58, two-tailed paired t-test, not significant).

**Supplementary Figure 4 –.**
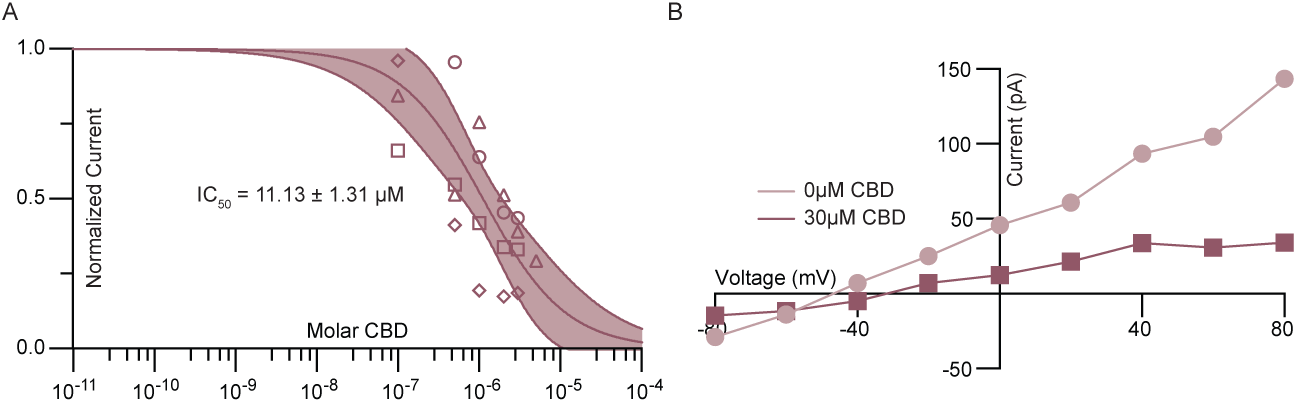
CBD inhibition of *Danio rerio* TREK-1. Dose-response curve. Data from individual patches are plotted with different shapes. Solid line is the fit to a four parameter logistic curve with 95% confidence interval shaded. IC_50_ = 11.1±1.3µM (mean±sem, n = 4 patches). **(b)** Representative current-voltage relationship from a *Danio rerio* TREK-1-containing patch before and after application of 30 µM CBD.

**Supplementary Figure 5 –.**
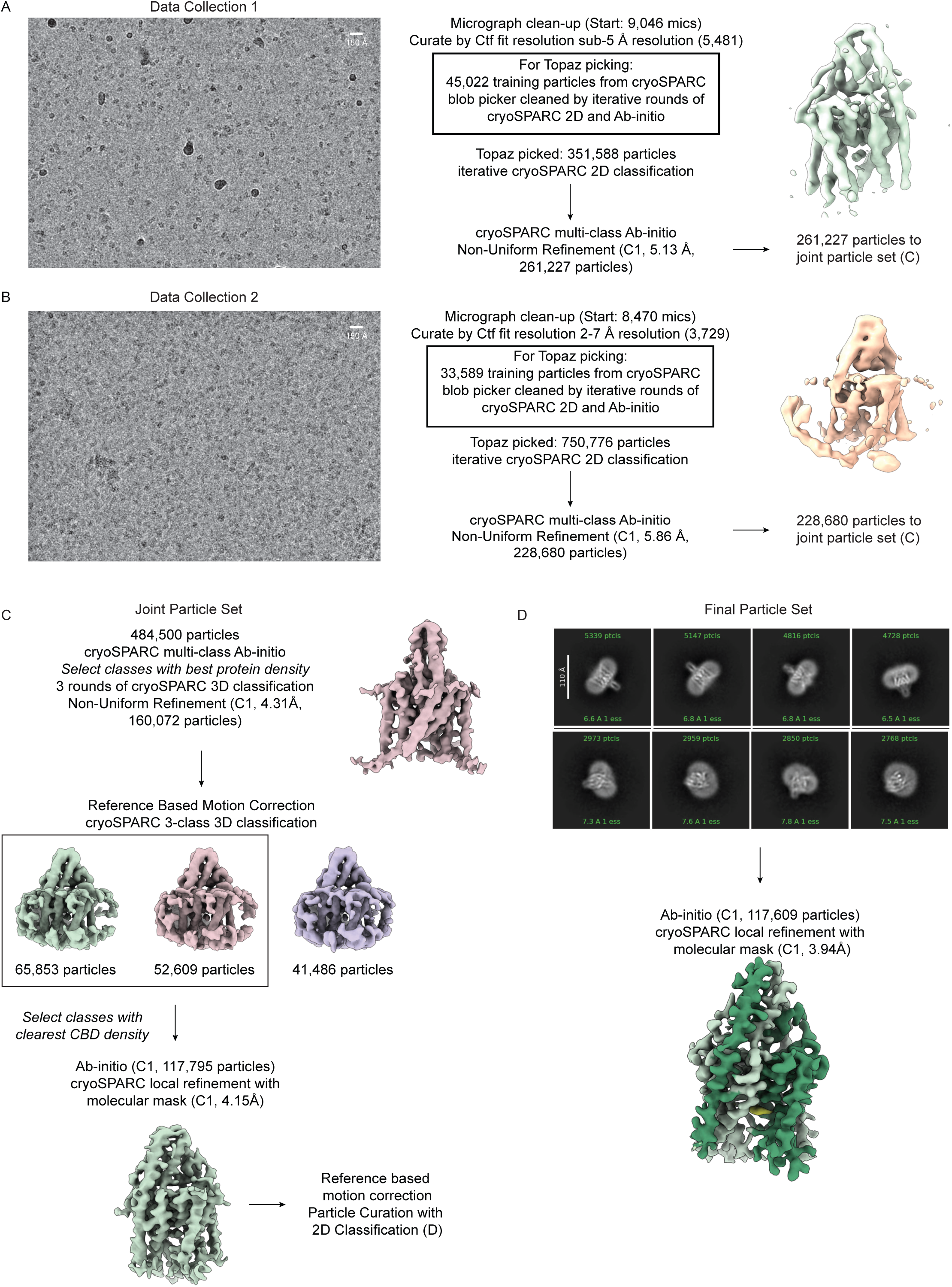
Cryo-EM data processing pipeline for TREK-1:CBD. **(a,b)** Representative micrograph and initial cryo-EM data processing from each dataset, **(c)** cryo-EM data processing pipeline from merged particle stack, and **(d)** representative 2D class averages and reconstructed map from final particle stack.

**Supplementary Figure 6 –.**
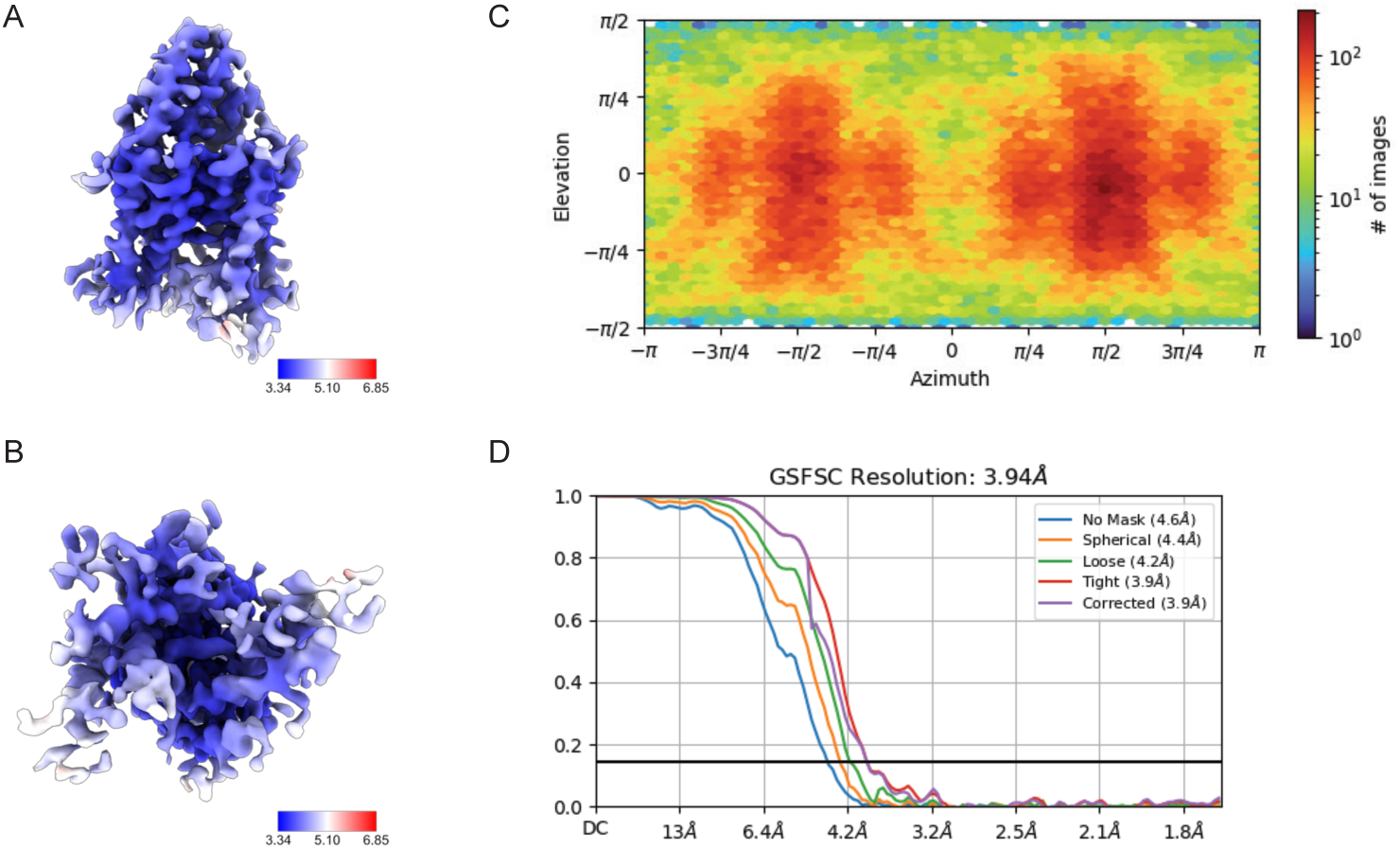
Cryo-EM structure validation for TREK-1:CBD. Final sharpened map colored by local resolution and viewed from **(a)** the membrane plane and **(b)** the cytoplasmic side. **(c)** View angle distribution of final particle stack. **(d)** Fourier shell correlation (FSC) versus resolution between half maps from final refinement.

**Supplementary Figure 7 –.**
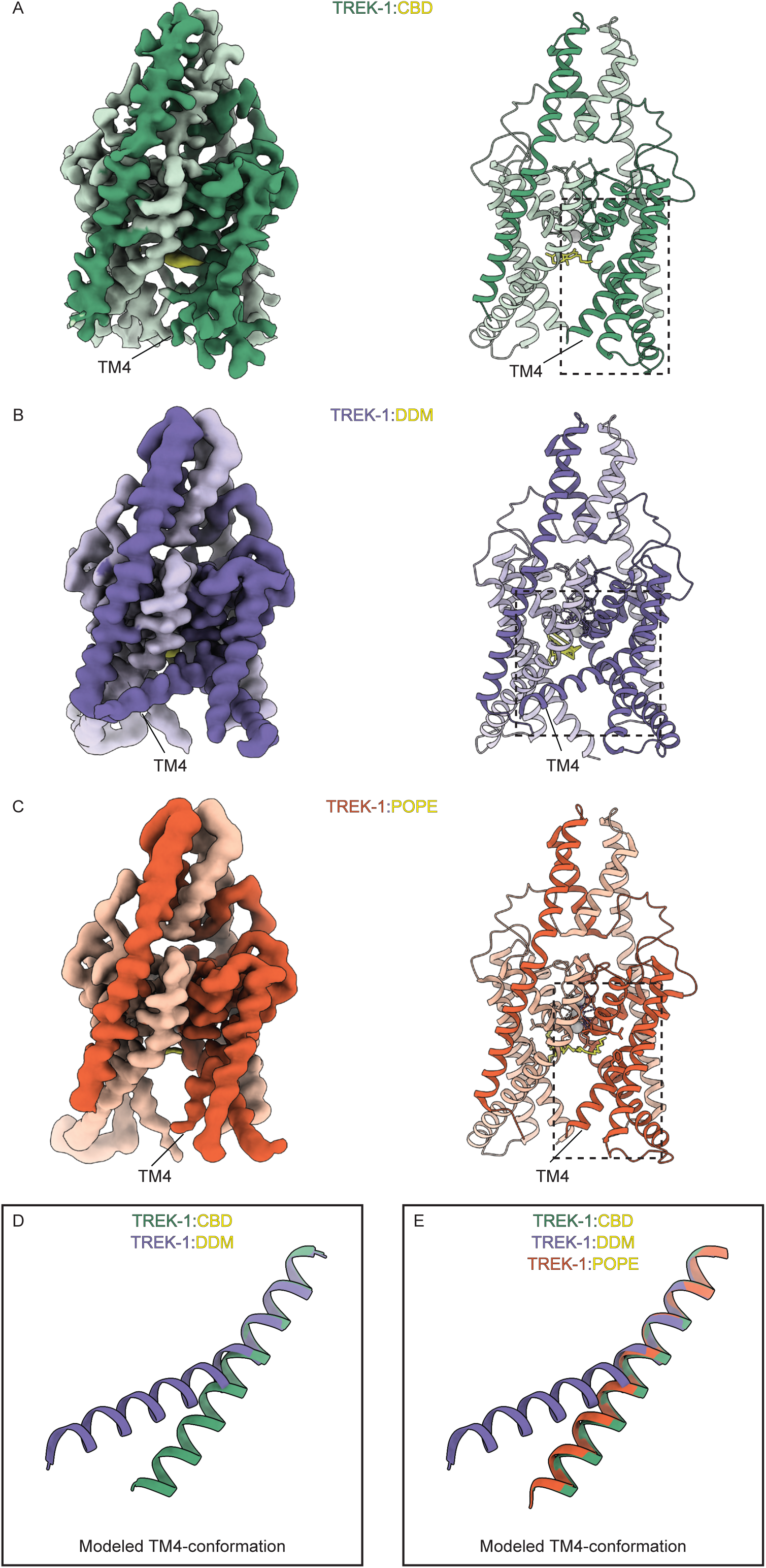
CBD-bound TREK-1 is stabilized in the TM4-down conformation. **(a)** Final map and structure of TREK-1:CBD focused on TM4. **(b)** Final map and structure of apo TREK-1 focused on TM4. **(c)** Final map and structure of TREK-1:POPE focused on TM4. **(d)** comparison of TM4 conformation in apo TREK-1 (purple) and TREK-1:CBD (green) and **(e)** TREK-1:POPE.

**Supplementary Figure 8 –.**
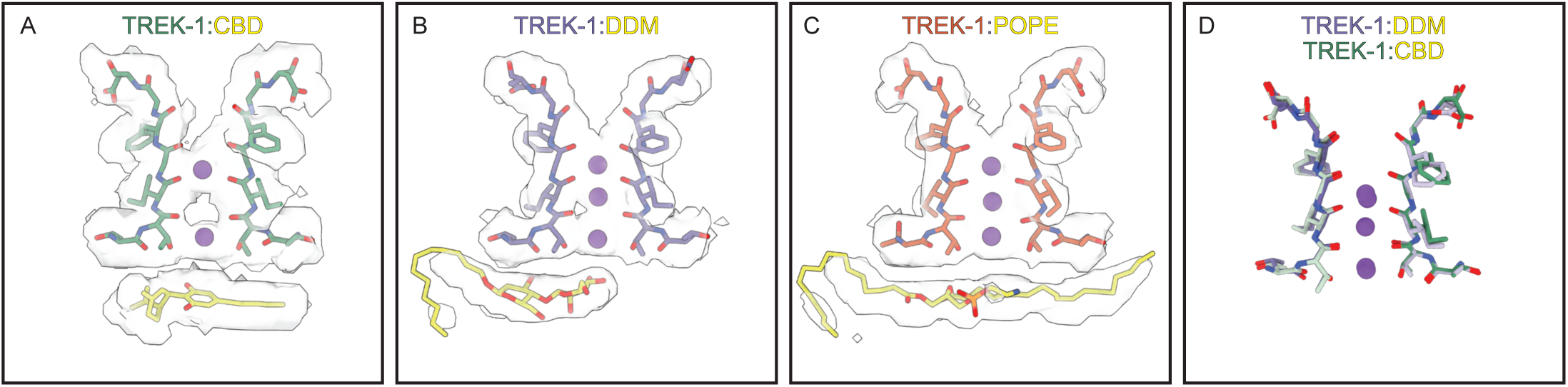
CBD-induced changes in the TREK-1 selectivity filter. Comparison of the TREK-1 selectivity filter and K^+^ density from **(a)** TREK-1:CBD, **(b)** apo TREK:1, and **(c)** TREK-1:POPE structures. **(d)** Overlaid selectivity filters from apo TREK-1 (purple) and TREK-1:CBD (green) structures.

**Supplementary Table 1 –.**
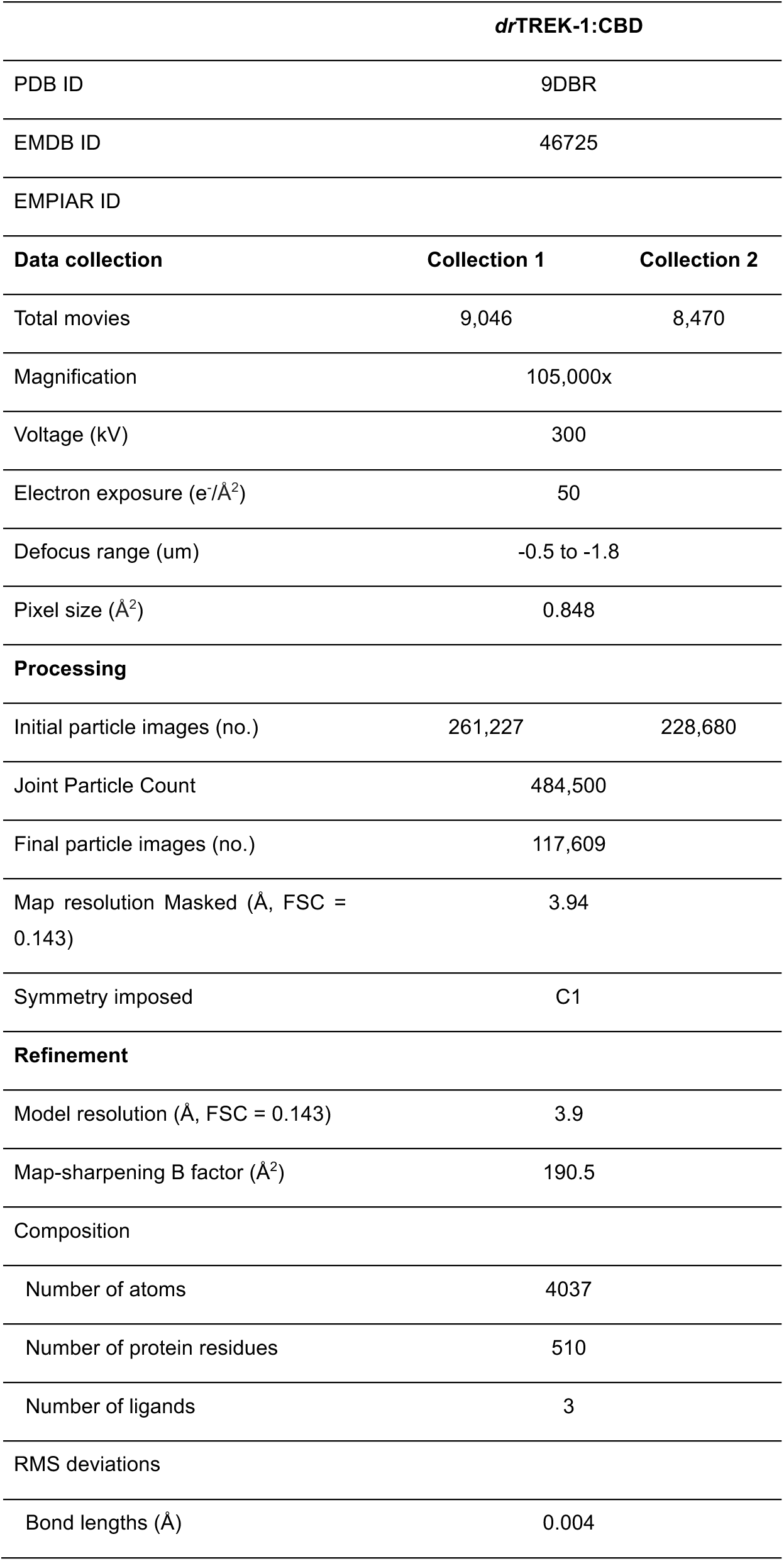
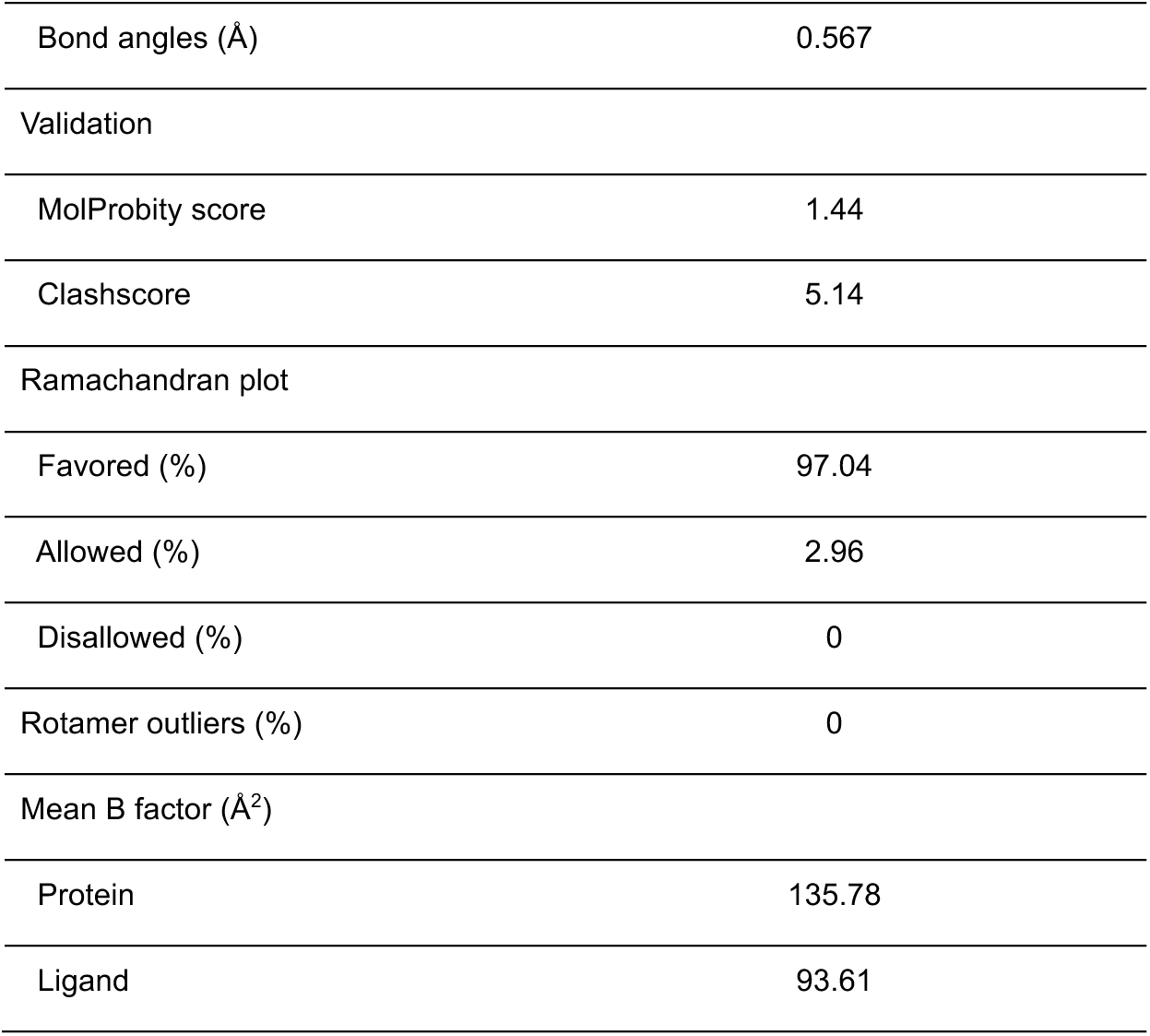
Cryo-EM data collection, refinement, and validation statistics.

| drTREK-1: CBD |  |  |
| --- | --- | --- |
| PDB ID | 9DBR |  |
| EMDB ID | 46725 |  |
| EMPIAR ID |  |  |
| Data collection | Collection 1 | Collection 2 |
| Total movies | 9,046 | 8,470 |
| Magnification | 105,000x |  |
| Voltage (kV) | 300 |  |
| Electron exposure (e<sup>-</sup>/Å<sup>2</sup>) | 50 |  |
| Defocus range (um) | -0.5 to -1.8 |  |
| Pixel size (Å<sup>2</sup>) | 0.848 |  |
| Processing |  |  |
| Initial particle images (no.) | 261,227 | 228,680 |
| Joint Particle Count | 484,500 |  |
| Final particle images (no.) | 117,609 |  |
| Map resolution Masked (Å, FSC = 0.143) | 3.94 |  |
| Symmetry imposed | C1 |  |
| Refinement |  |  |
| Model resolution (Å, FSC = 0.143) | 3.9 |  |
| Map-sharpening B factor (Å<sup>2</sup>) | 190.5 |  |
| Composition |  |  |
| Number of atoms | 4037 |  |
| Number of protein residues | 510 |  |
| Number of ligands | 3 |  |
| RMS deviations |  |  |
| Bond lengths (Å) | 0.004 |  |

|  |  |
| --- | --- |
| Bond angles (Å) | 0.567 |
| Validation |  |
| MolProbity score | 1.44 |
| Clashscore | 5.14 |
| Ramachandran plot |  |
| Favored (%) | 97.04 |
| Allowed (%) | 2.96 |
| Disallowed (%) | 0 |
| Rotamer outliers (%) | 0 |
| Mean B factor (Å<sup>2</sup>) |  |
| Protein | 135.78 |
| Ligand | 93.61 |

